# Altered resting-state brain entropy (BEN) by rTMS across the human cortex

**DOI:** 10.1101/2024.07.16.601109

**Authors:** Dong-Hui Song, Xin-Ping Deng, Yuan-Qi Shang, Da Chang, Ze Wang

## Abstract

Repetitive transcranial magnetic stimulation (rTMS) is a non-invasive brain stimulation method effective in treating various neuropsychiatric disorders, yet its mechanisms are not fully understood. In general, rTMS protocols are categorized into excitatory protocols including high-frequency rTMS (HF-rTMS) and intermittent theta burst stimulation (iTBS), and inhibitory protocols including low-frequency rTMS (LF-rTMS) and continuous theta burst stimulation (cTBS). Brain entropy (BEN) measures irregularity, disorders, and complexity of brain activity, our previous studies have indicated that BEN affects excitatory rTMS, including HF-rTMS and iTBS. However, two important questions remain whether rTMS is equally sensitive to inhibitory rTMS and whether it can induce opposite brain activities, another question concerns whether rTMS can induce specific changes across brain regions. To address these issues, we utilized our own cTBS targeted on the left dorsal lateral prefrontal cortex (L-DLPFC) dataset and publicly available LF-rTMS dataset with stimulating sites including the L-DLPFC, left temporal parietal junction (L-TPJ), and left occipital cortex (L-OCC), from the OpenNeuro. BEN maps were calculated before and after stimulation. The results showed that L-DLPFC cTBS increased BEN in the MOFC and L-DLPFC LF-rTMS increased BEN in the MOFC, subgenual anterior cingulate cortex (MOFC/sgACC) and putamen, the regions are consistent with our previous findings with HF-rTMS and iTBS. Additionally, L-TPJ LF-rTMS resulted in increased BEN in the right TPJ, while L-OCC LF-rTMS led to decreased BEN in the posterior cingulate cortex (PCC). Our findings suggest that BEN is not only sensitive to excitatory rTMS but also to inhibitory rTMS. Moreover, LF-rTMS induces different effects across brain regions, as detected by BEN.

## 1. Introduction

Transcranial magnetic stimulation (TMS) is the most widely utilized method to non-invasively modulate human brain activity and investigate the causal relationship between brain function and behavior (Machii, Cohen et al. 2006). It is also being explored as a potential treatment for psychiatric disorders such as major depressive disorder (MDD), Alzheimer’s disease (AD), addictive disorders, and attention deficit hyperactivity disorder (ADHD) (Ridding and Rothwell 2007, Gorelick, Zangen et al. 2014, Chang, Zhang et al. 2018, Lefaucheur, Aleman et al. 2020, Antonelli, Fattore et al. 2021, Somaa, de Graaf et al. 2022). It has been shown through various studies that TMS can induce neuroplasticity across brain regions and time scales, linked to long-term potentiation and long-term inhibition. TMS protocols are categorized into excitatory protocols, such as conventional high-frequency rTMS (HF-rTMS) and intermittent theta burst stimulation (iTBS), and inhibitory protocols, such as conventional low-frequency rTMS (LF-rTMS) and continuous theta burst stimulation (cTBS) based on their impact on the motor cortex. Although inhibitory effects are later generalized from the motor cortex to other cortical regions, an increasing body of research suggests that this inhibitory effect established in the motor cortex may not generalize to other cortical regions (Eldaief, Halko et al. 2011, Caparelli, Backus et al. 2012, Mancini, Mastropasqua et al. 2017, Castrillon, Sollmann et al. 2020). A recent study by Castrillon et al (2020) reported that LF-rTMS had the opposite impact on the functional connectivity (FC) of sensory (occipital cortex, OCC) and cognitive brain regions (left dorsal-lateral prefrontal cortex, L-DLPFC). Using spectral dynamic causal modeling (DCM), the authors identified that stimulation of the L-DLPFC decreased local inhibition and disrupted feedforward and feedback connections; identical stimulation increased local inhibition and enhanced forward signaling in the OCC (Castrillon, Sollmann et al. 2020). Some commonly utilized resting-state measurement metrics, including the amplitude of low-frequency fluctuation (ALFF) (Yang, Long et al. 2007, Zou, Zhu et al. 2008), regional homogeneity (ReHo) (Zang, Jiang et al. 2004), and signal variability (McGonigle, Howseman et al. 2000) also were analyzed in the study. Interestingly, none of these metrics revealed variances in the stimulated regions, despite the fascinating post-stimulation effects observed off the target from FC and DCM. It’s possible that the stimulation does not directly cause changes in local brain activity, but rather affects distant areas of the brain. A recent review indicated that rTMS don’t necessarily cause brain activity effects in the immediate stimulation sites of the cortex (Rafiei and Rahnev 2022). Furthermore, it is plausible that the selected resting state indicators may lack the requisite sensitivity to detect post-stimulation effects. The purpose of this study was to re-examine the regional effects of LF-rTMS using an emerging resting state fMRI (rs-fMRI) metric: brain entropy (BEN) (Wang, Li et al. 2014, Wang 2021).

The concept of entropy originates from thermodynamics (Boltzmann 1974) and information theory (Shannon 1948), measuring the disorder and unpredictability of a system, often used to gauge the complexity of a dynamic system (Pincus 1991). In recent years, there has been a growing interest in the nonlinear dynamic measurements of rs-fMRI signals (Friston, Harrison et al. 2003, Smith, Yan et al. 2014, Wang, Li et al. 2014, Song, Jann et al. 2024). Among these methods, BEN has emerged as a promising approach that is associated with multiple cognitive abilities (Wang 2021, Del Mauro and Wang 2024), task performance (Lin, Chang et al. 2022, Camargo, Del Mauro et al. 2024, Song and Wang 2024), hormones (Song and Wang 2024), and neuropsychiatric disorders (Zhou, Zhuang et al. 2016, Xue, Yu et al. 2019, Liu, Song et al. 2020, Wang and Initiative 2020, Jiang, Cai et al. 2023, Dong-Hui Song 2024). As well as BEN is sensitive to rTMS effects. We observed that HF-rTMS over L-DLPFC reduced BEN in the medial orbitofrontal cortex and subgenual anterior cingulate cortex (MOFC/sgACC) (Song, Chang et al. 2019) and subthreshold and suprathreshold iTBS-induced converse BEN in the striatum in healthy young adults (Liu, Song et al. 2024). BEN also reflects patterns related to depression and its treatment (Lin, Lee et al. 2019, Liu, Song et al. 2020, Dong-Hui Song 2024). Furthermore, BEN was shown to provide complementary information to fractional ALFF (Zou, Zhu et al. 2008) and cerebral blood flow (CBF) (Wang, Aguirre et al. 2008, Li, Zhu et al. 2012, Song, Chang et al. 2019). The subsequent scientific questions are whether BEN is similarly sensitive to the effects of LF-rTMS and cTBS and whether BEN is sensitive to the effects of different target regions. To address the two questions, we utilized our own collected cTBS dataset and LF-rTMS dataset from the OpenNeuro ds001927 dataset released by Castrillon et al (2020).

## 2. Methods

### 2.1 The cTBS dataset

#### 2.1.1 Participants

Eighteen healthy young adults (mean age 22.67± 3.25 years, 9 males) who were right-handed were recruited from Hangzhou Normal University and the local community. All participants had no history of psychiatric illness or neurological disorders and were instructed to refrain from caffeine, alcohol, nicotine, or any drugs for at least 24 hours before the start of the experiment. The study was approved by the local internal review board at Hangzhou Normal University and adhered to the Declaration of Helsinki, written informed consent was acquired from each participant.

#### 2.1.2 Experimental Procedure

Each participant received two sessions of MRI on two separate days 48 hours apart with the cTBS or without cTBS using the same imaging protocols. The order of stimulation was randomized and counterbalanced across participants. On the first day, each participant underwent a T1-weighted anatomical MRI scan to localize the stimulation targets and determine the resting motor threshold (rMT). As the stimulation order was counterbalanced, half of the participants received cTBS on the first day, while the other half received cTBS on the second day. The post-cTBS scan was performed within 15 min to ensure that the stimulation effects were measured.

#### 2.1.3 Brain stimulation

The cTBS was performed using a Magstim Rapid stimulator with a figure-of-eight coil (Magstim Ltd, Whitland, UK). Before stimulation began, participants were instructed to lie down in a chair with a headrest to support their heads, and the stimulation target and resting motor threshold (rMT) were determined. The stimulation location was targeted using the BrainSight navigation system (Rogue Research Inc) which can dynamically visualize the TMS coil in 3D space in real-time position on top of the individual structural image. The coordinate of the target site in the left DLPFC was set to be [−40, 26, 37] in MNI space. The rMT intensity was determined for each participant by stimulating the left primary motor cortex (M1) with single pulses and detecting corresponding muscle twitching of the relaxed contralateral first dorsal interosseous (FDI) muscle. For cTBS, the coil was positioned tangentially to the skull surface above the left DLFPC with the handle pointed backward at a 45° angle. The cTBS protocol with three pulses of stimulation given at 50 Hz, repeated every 200 ms with an intensity of 80% of the rMT. A total of 600 pulses were administrated which equals 40s of stimulation. The parameters and procedures of cTBS can also be found in (Shang, Chang et al. 2020).

#### 2.1.4 MRI acquisition

The images were acquired on a GE Discovery MR 750 3Tesla scanner (General Electric, Waukesha, WI, USA) at the Center for Cognition and Brain Disorders at Hangzhou Normal University, China. During the scanning process, a comfortable and tight cushion was placed to immobilize the head and reduce motion. The participants were instructed to relax and remain still with their eyes closed, not to fall asleep.

High-resolution T1-weighted anatomical images were acquired with a 3D spoiled gradient echo sequence (3D-SPGR) with repetition time/echo time (TR/TE) = 8.1/3.39 ms, flip angle (FA) = 7°, field of view (FOV) = 256 × 256 mm^2^, matrix = 256 × 256, 1.0 mm^3^ isotropic voxels, 176 slices without interslice gap. The functional images were acquired with a T2*-weighted gradient-echo EPI pulse sequence with the following parameters: TR/TE = 2000/30 ms, flip angle = 90°, field of view = 220 × 220 mm^2^, matrix = 64 × 64, 3.4 mm^3^ isotropic spatial resolution, and 37 interleaved slices. Each fMRI run lasted for 6 minutes, during which 180 functional volumes were acquired.

#### 2.1.5 MRI preprocessing

MRI preprocessing was performed using Statistical Parametric Mapping (SPM12, WELLCOME TRUST CENTRE FOR NEUROIMAGING, London, UK, http://www.fil.ion.ucl.ac.uk/spm/software/spm12/) (Friston, Holmes et al. 1994). Structural images were segmented into gray matter (GM), white matter (WM), and cerebrospinal fluid (CSF) and registered into the Montreal Neurological Institute (MNI) standard brain space.

The following steps were used for the functional images. 1) The first 6 volumes were discarded to allow image intensity to reach a steady state. 2) The remaining images were corrected for slice timing acquisition difference using the middle slice as the reference and then corrected for head motions using the first image volume as the reference. 3) The functional images were spatially registered with the high-resolution structural images. WM and CSF segmentation maps were back registered into the functional images and resampled to the same resolution for extracting average WM and CSF signals; 4) Denoising was performed via simple regression by including six head motion parameters, WM signal, and CSF signal. 5) The functional images were smoothed with an isotropic Gaussian kernel with a full-width-at-half-maximum (FWHM) of 6 mm^3^; 6) The band-pass filtered (0.01–0.08 Hz) were performed to eliminate high-frequency noise and low-frequency drift; 7) The functional images were warped into the MNI space using the spatial transform estimated from the structural images, and resampled with a resolution of 3[×[3[×[3 mm^3^.

### 2.2 The LF-rTMS dataset

#### 2.2.1 Participants

The dataset from OpenNeuro ds001927 (https://openneuro.org/datasets/ds001927/versions/2.1.0) comprised twenty-three healthy young adults (mean age = 25.74 ± 3.22 years, 12 females) with right-handed and without any psychiatric condition. The original study was approved by the local institutional review board and was conducted by the Declaration of Helsinki.

#### 2.2.2 Brain stimulation

LF-rTMS was performed using an electric-field–navigated Nexstim eXimia system (version 4.3; Nexstim Plc, Helsinki, Finland) and a biphasic figure-of-eight stimulation coil. Before any rTMS session, the participant’s head was co-registering to their structural image by the neuronavigation system, allowing it to continuously track the coil position to the individual target region via infrared cameras. The individual rMT was determined by the maximum likelihood algorithm by mapping the cortical representation of the right abductor pollicis brevis muscle using surface muscle electrodes (Neuroline 720; Ambu, Ballerup, Denmark) and an integrated electromyography device. During stimulation, the coil was positioned perpendicular to the skull surface, LF-rTMS with a frequency of 1 Hz and a stimulation intensity of 100% of the individual rMT (mean rMT = 34.4%, SD = 7.5%) was applied for 20 minutes to three target regions: L-DLPFC (mean MNI coordinate =[-48,43,35]), left OCC (L-OCC) (mean MNI coordinate=[-14,-117,-13]), and left temporoparietal junction (L-TPJ) (mean MNI coordinate=[-77,-60,13]), with a total of 1200 rTMS pulses. The parameters and procedures of LF-rTMS can also be found in (Castrillon, Sollmann et al. 2020).

#### 2.2.3 MRI acquisition

MRI data were acquired on a 3T Philips Ingenia MRI scanner using the body coil for transmission and the 32-channel head coil for signal reception (Philips Healthcare, Best, The Netherlands). During the scanning process, the scanner room was dimmed, and participants were instructed to remain awake with their eyes open.

High-resolution T1-weighted anatomical images were acquired with a 3D T1-weighted turbo field echo (3D-TFE) with TR/TE = 9/3.98 ms, FA= 8°, FOV = 256 × 256 mm^2^, matrix = 256 × 256, 1.0 mm^3^ isotropic voxels, 170 slices without interslice gap. The functional images were acquired with multiband with the following parameters: multiband factor = 2; SENSE factor = 2; TR/TE = 1250/30ms; FA = 70°; FOV = 192 × 192 mm^2^; matrix size = 64 × 64; voxel size = 3 × 3 × 3 mm^3^). Each fMRI run lasted for 12.35 minutes, during which 600 functional volumes were acquired.

#### 2.2.4 MRI preprocessing

The MRI preprocessing was performed using fmriprep (Esteban, Markiewicz et al. 2019) (version=23.1.4) (https://pypi.org/project/fmriprep-docker/), which is based on Nipype (version=1.8.6) (Gorgolewski, Burns et al. 2011), is containerized to docker (version=2 4.0.7) (https://www.docker.com/), then denoising using based Nilearn (version=0.10.2) (https://nilearn.github.io/stable/index.html) customed Python (version=3.10) (https://www.python.org/) scripts.

The structural images were skull-stripped and segmented into GM, WM, and CSF using fast (Zhang, Brady et al. 2001), then normalized to MNI space. The functional images underwent preprocessing through the following steps: (1) Head motion correction was performed using mcflirt (Jenkinson, Bannister et al. 2002). (2) The functional reference was then co-registered to the T1w reference using mri_coreg followed by flirt (Jenkinson and Smith 2001), framewise displacement (FD) (Power, Barnes et al. 2012) and DVARS were calculated. The average signals were extracted within the GM, CSF, and WM. (3) The functional images were resampled into MNI space using antsApplyTransforms (ANTs). (4) Detrending and temporal bandpass filtering (0.009-0.08 Hz) was performed. (5) Nuisance components were regressed out including the six motion parameters, FD, std_dvars, rmsd, tcompcor, and first three componence of c_comp_cor, as well as the average WM signals and average CSF signals. (6) Smoothing with an isotropic Gaussian kernel (FWHM = 6 mm). For more detailed preprocessing details, please refer to (Liu, Song et al. 2024).

### 2.3 BEN mapping calculation

The voxel-wise BEN maps were calculated from the preprocessed rs-fMRI using the BEN mapping toolbox (BENtbx) (Wang, Li et al. 2014) based on sample entropy (Richman and Moorman 2000). The toolbox can be found at https://www.cfn.upenn.edu/zewang/BENtbx.php and https://github.com/zewangnew/BENtbx. The window length was set to 3 and the cutoff threshold was set to 0.6 according to our experimental optimization (Wang, Li et al. 2014). The first four volumes were discarded for signal stability, BEN maps were smoothed with an isotropic Gaussian kernel with FWHM = 8 mm. More details of BEN calculation can be found in our previous studies (Wang, Li et al. 2014, Song, Chang et al. 2019, Song, Chang et al. 2019, Wang and Initiative 2020, Wang 2021, Lin, Chang et al. 2022).

### 2.4 Statistical analysis

Paired sample T-tests were performed to determine the difference in BEN between pre- and post-stimulus conditions, statistical significance was set to voxel-wise p < 0.001 and cluster size > 135 mm^3^.

## 3. Results

The cTBS over the L-DLPFC increased BEN in the MOFC (see Fig 1), while LF-rTMS over L-DLPFC similarly increased BEN in the MOFC but also involved the sgACC. LF-rTMS over L-OCC decreased BEN in the posterior cingulate cortex (PCC), while LF-rTMS over L-TPJ increased BEN in the right posterior middle temporal gyrus and TPJ (pMTG/TPJ) (Fig 2). Table 1 showed information on clusters exceeding the threshold (voxel-wise p < 0.001, cluster size ≥ 135 mm^3^), including peak MNI coordinates, cluster size, label on the AAL atlas, and brain regions.

**Fig 1.**
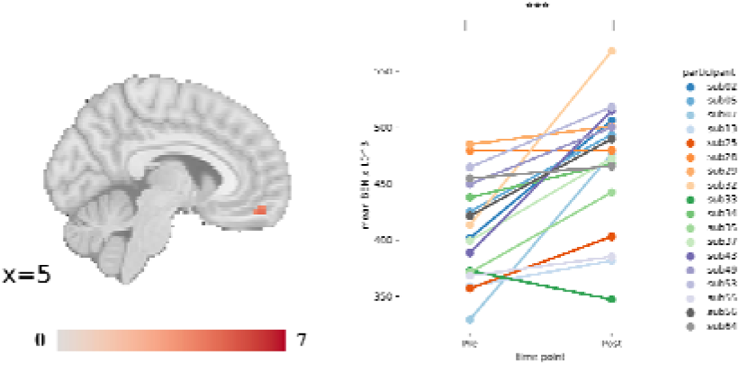
Increased BEN by iTBS over L-DLPFC. The left represents the increased BEN; the right represents the mean BEN changes for each participant from the clusters with significant BEN changes by iTBS over L-DLPFC. The number beneath each slice indicates its location along the x-axis in the MNI space, the colobar indicates t values, and the red indicates higher BEN after iTBS. The y-axis represents the mean BEN values × 10^^3^ within the cluster, while the x-axis corresponds to different time points (Pre vs Post) of iTBS.

**Fig 2.**
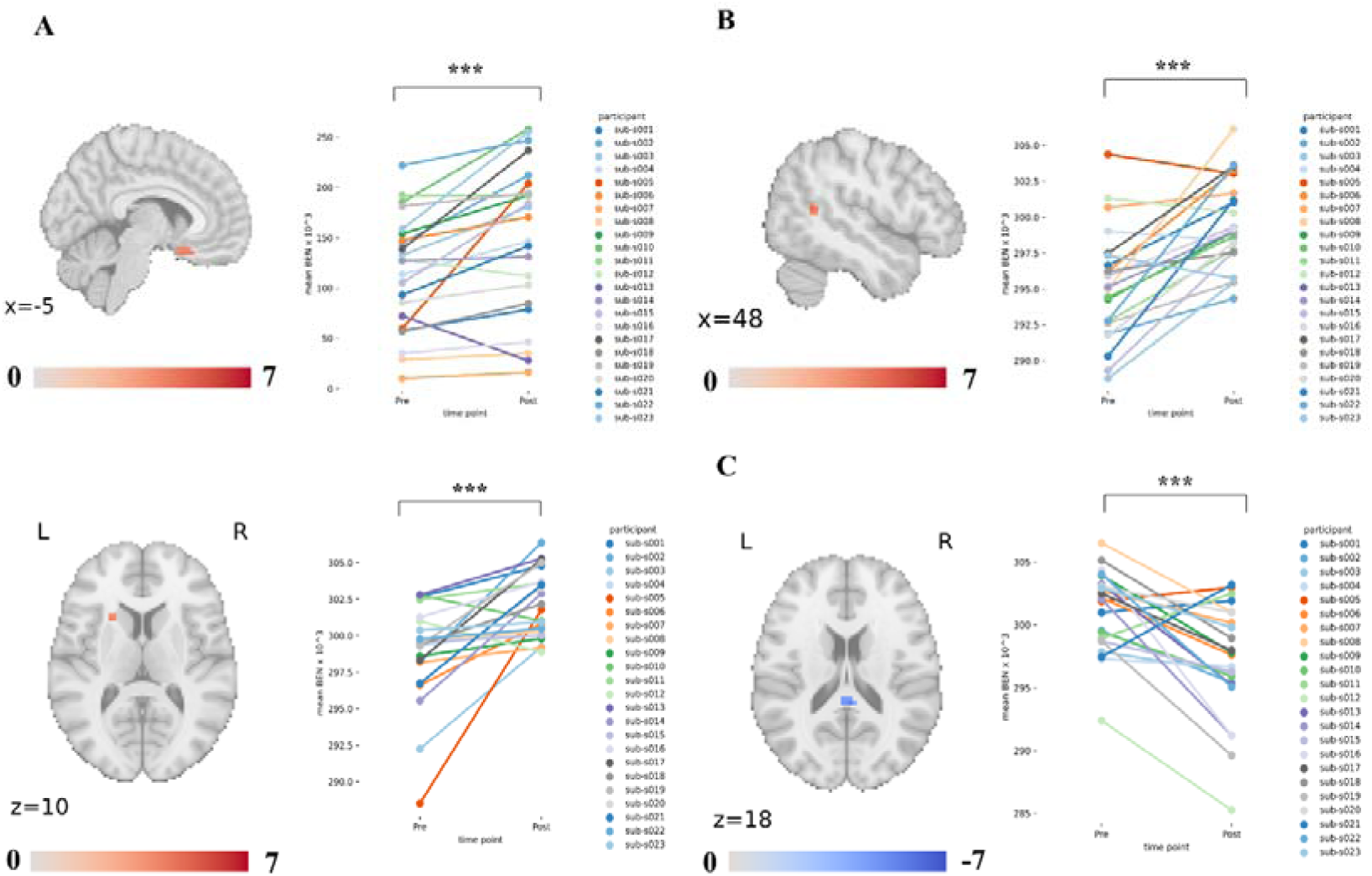
BEN was altered by LF-rTMS across the human cortex. A) The left represents the increased BEN; the right represents the mean BEN changes for each participant from the clusters with significant BEN changes by LF-rTMS over L-DLPFC. B) The left represents the increased BEN; the right represents the mean BEN changes for each participant from the cluster with significant BEN changes by LF-rTMS over L-TPJ. C) The left represents the decreased BEN; the left represents the mean BEN changes for each participant from the cluster with significant BEN changes by LF-rTMS over L-OCC. The number beneath each slice indicates its location along the x or z-axis in the MNI space, the colobar indicates t values, the red indicates higher BEN after rTMS, and the blue indicates lower BEN after rTMS. The y-axis represents the mean BEN values × 10^3 within the cluster, while the x-axis corresponds to different time points (Pre vs Post) of rTMS.

**Table 1.** Clusters size table for BEN analysis.

| Contrast<br>(Post-Pre) | Cluster<br>ID | Peak MNI<br>Coordinate (X,<br>Y, Z) | Peak<br>Value<br>(t) | Cluster<br>Size<br>(mm<sup>3</sup>) | AAL Label | Brain Region |
| --- | --- | --- | --- | --- | --- | --- |
| L-DLPFC cTBS | 1 | (9,54, -18) | 4.86 | 324 | Rectus_R | MOFC |
| L-DLPFC<br>LF-rTMS | 1 | (-15,15, -18) | 5.36 | 3672 | Frontal_Sup_Orb_L | MOFC/sgACC |
|  | 1a | (-27,18, -27) | 4.57 |  |  |  |
|  | 2 | (-21,15,12) | 4.28 | 216 | Putamen_L | Putamen |
| L-TPJ LF-rTMS | 1 | (48, -54,12) | 4.84 | 162 | Temporal_Mid_R | pMTG/TPJ |
| L-OCC LF-rTMS | 1 | (3, -39, 18) | -5.30 | 486 | Cingulum_Post_R | PCC |

## 4. Discussion

Our results indicated that rTMS across brain regions leads to distinct patterns of BEN changes. Specifically, the cTBS and LF-rTMS over L-DLPFC resulted in increased BEN in the MOFC/sgACC, and BEN of left putamen was increased by LF-rTMS over L-DLPFC. The LF-rTMS over L-OCC led to decreased BEN in the PCC, and LF-rTMS over L-TPJ caused increased BEN in the right pMTG/TPJ.

The cTBS and LF-rTMS have been widely demonstrated to have similar effects on decreasing cortical excitability (Chen, Classen et al. 1997, Muellbacher, Ziemann et al. 2000, Huang, Edwards et al. 2005, Wischnewski and Schutter 2015, Suppa, Huang et al. 2016), while they have opposite effects compared to HF-rTMS and iTBS that increase cortical excitability (Speer, Kimbrell et al. 2000, Huang, Edwards et al. 2005, Jung, Shin et al. 2008, Guse, Falkai et al. 2010, Huang, Rothwell et al. 2011). In the study, the cTBS over L-DLPFC and LF-rTMS over L-DLPFC also demonstrated similar effects in the MOFC/sgACC, although the effect region of cTBS appears to be more prominent. This indicates that both rTMS protocols act on the MOFC/sgACC and result in increased BEN. The region aligns with the discovery from our previous studies on HF-rTMS. Our previous studies using HF-rTMS (20 Hz) showed lower BEN or higher temporal coherence (Song, Chang et al. 2019, Wang 2021, Song Donghui 2023) and higher fractional ALFF in the MOFC/sgACC (Xue, Guo et al. 2017). A negative correlation between BEN and fALFF was also demonstrated in the MOFC (Song, Chang et al. 2019). In contrast to the results of HF-rTMS over L-DLPFC, which resulted in decreased BEN, increased BEN by LF-rTMS or cTBS over L-DLPFC means increased brain activity irregularity and decreased long-range temporal coherence (LRTC) in MOFC/sgACC (Wang 2021). The MOFC/sgACC plays a pivotal role in emotional regulation (Kim and Hamann 2007), reward processing (Rolls 2000), decision-making (Bechara, Damasio et al. 2000, Fellows 2007), and inhibition control (Elliott and Deakin 2005), functions frequently disrupted in psychiatric disorders, such as MDD (Liu, Song et al. 2020) and addictive disorders (Volkow and Fowler 2000, Koob 2009). The beneficial effects of HF-rTMS over the L-DLPFC have been widely reported in the literature. These beneficial effects stem from the top-down regulation of the L-DLPFC over the MOFC/sgACC. In contrast, LF-rTMS or cTBS over the L-DLPFC has inversely results (Kimbrell, Little et al. 1999, Skrdlantova, Horacek et al. 2005, Speer, Benson et al. 2009) that may be attributed to the reduction of the L-DLPFC’s top-down regulation over the MOFC/sgACC by LF-rTMS or cTBS over the L-DLPFC. We also observed increased BEN in the left putamen when applying LF-rTMS to the L-DLPFC. In contrast, our previous study on intermittent theta burst stimulation (iTBS) over L-DLPFC showed that subthreshold iTBS resulted in decreased BEN. At the same time, suprathreshold iTBS led to increased BEN in the putamen (Liu, Song et al. 2024). The fronto-striatal circuit in modulating reward processing and cognitive control by top-down regulation has been demonstrated in numerous studies (Chudasama and Robbins 2006, Staudinger, Erk et al. 2011, Morris, Kundu et al. 2016, Becker, Kirsch et al. 2017), increased BEN in putamen may disrupt the regulatory effects on the frontostriatal circuit.

The LF-rTMS over L-TPJ enhanced BEN in the right pMTG/TPJ. The pMTG/TPJ is key in social cognition (Krall, Rottschy et al. 2015, Sowden and Catmur 2015, Wilterson, Nastase et al. 2021). A study indicated that LF-rTMS over L-TPJ reduces hostile attributions (Giardina, Caltagirone et al. 2011). Moreover, LF-rTMS over L-TPJ is widely used in treating auditory verbal hallucinations (AVH) in schizophrenia and is believed to effectively reduce AVH symptoms by decreasing the FC of both left and right rTPJ. This reduction in FC aligns with increased activation in the right TPJ, primarily due to heightened right hemisphere activity, resulting in greater temporal dynamics. Conversely, there is no significant change in activation in the left TPJ, leading to desynchronization in the time series of these two regions. The heightened activity in the right TPJ diminishes the interaction between left and right TPJ, consequently reducing AVH symptoms. LF-rTMS over the L-OCC reduced BEN of the PCC, the results align with the original study that reported an increase in FC between L-OCC and PCC (Castrillon, Sollmann et al. 2020).

Our study indicates that BEN remains sensitive to various rTMS protocols, including stimulation frequency and targets. Furthermore, LF-rTMS and cTBS applied over the L-DLPFC demonstrate similar effects, but divergently affect the MOFC/sgACC compared to HF-rTMS. Thus, excitatory, and inhibitory rTMS evoke opposing effects in remote brain regions when targeting the L-DLPFC.

## Acknowledgments

We thank Castrillon et al for releasing their dataset.

## Data and code availability

The cTBS dataset is available in the online repository of OpenNeuro ds001927 (https://openneuro.org/datasets/ds001927/versions/2.1.0).

fmriprep is available at https://fmriprep.org/en/0.6.4/index.html.

Nilearn is available at https://nilearn.github.io/dev/index.html.

SPM is available at https://www.fil.ion.ucl.ac.uk/spm/software/spm12/.

Further updates related to this study will be available at https://github.com/donghui1119/Brain_Entropy_Project/tree/main/NIBS/LF-rTMS (upon publication of the manuscript).

## CRediT authorship contribution statement

Dong-Hui Song: conceptualization, data curation, data analysis, visualization, manuscript drafting, and editing. Xin-Ping Deng: manuscript editing, Yuan-Qi Shang: data curation. Da Chang: data curation. Ze Wang: conceptualization, manuscript editing and final manuscript proofing, supervision, project administration.

